# Modelling altered signalling of G-protein coupled receptors in inflamed environment to advance drug design

**DOI:** 10.1101/2021.10.20.465073

**Authors:** Sourav Ray, Arne Thies, Vikram Sunkara, Hanna Wulkow, M. Özgür Celik, Fatih Yergöz, Christof Schütte, Christoph Stein, Marcus Weber, Stefanie Winkelmann

## Abstract

We previously reported the successful design, synthesis and testing of the prototype opioid painkiller NFEPP that does not elicit adverse side effects. Uniquely, this design was based on mathematical modelling of extracellular interactions between G-protein coupled receptors (GPCRs) and ligands, recognizing that GPCRs function differently under pathological versus healthy conditions. We now present a novel stochastic model of GPCR function that includes intracellular dissociation of G-protein subunits and modulation of plasma membrane calcium channels associated with parameters of inflamed and healthy tissue (pH, radicals). The model is validated against *in vitro* experimental data for NFEPP and fentanyl ligands at different pH values. We found markedly reduced calcium channel inhibition induced by NFEPP at normal pH compared to lower pH, in contrast to the effect of fentanyl, and enhanced constitutive G-protein activation but lower probability of ligand binding with increasing radical concentrations. By means of molecular dynamics simulations, we also assessed qualitative changes of reaction rates due to additional disulfide bridges inside the GPCR binding pocket. The results suggest that, compared to radicals, low pH is a more important determinant of overall GPCR function in an inflamed environment. Future drug design efforts should take this into account.

## 1 Introduction

The family of G-protein coupled receptors (GPCRs) represents the largest class of receptors in the human genome and some of the most common drug targets. Located on the cell membrane, they transduce extracellular signals into key physiological effects. Natural GPCR ligands include neurotransmitters, chemokines, hormones, odours or photons. GPCRs are involved in a large number of disorders, such as diabetes, high blood pressure, depression, addiction, pain, arthritis, Parkinson’s and many others [1]. A prominent member of this family is the *µ*-opioid receptor (MOR).

Recent works of our group [2] led to the development of the novel analgesic compound *N*-(3-fluoro-1-phenethylpiperidin-4-yl)-*N*-phenylpropionamide (NFEPP) which activates the MOR preferentially at acidic extracellular pH-levels, as given in injured tissues [3]. This is of utmost interest because it may preclude the adverse effects of conventional MOR agonists like fentanyl which include constipation, sedation and apnea. These adverse effects are mediated mostly in the brain and the gut, i.e. in healthy tissues (pH 7.4). Since the generation of pain can be effectively inhibited by blocking the electrical excitation of sensory neurons at the site of the injury (i.e. the origin of nociceptive stimulation), this gives rise to the hope that NFEPP might have less or even no adverse effects, which could already be corroborated in animal studies [2, 4, 5, 6].

Up to now, the effects of NFEPP and fentanyl were mathematically analysed at the level of their binding rates at relevant amino acid residues accessible from the extracellular side of MOR [2, 7]. To get a more complete picture, we herein present a model of the intracellular second messenger pathways relevant to pain and analgesia. The mechanism underlying the analgesic effect of MOR activation in nociceptive neurons is mainly due to a stabilisation or even lowering of the plasma membrane potential beneath the threshold value required to elicit an action potential (reviewed in [3, 8]). This effect is mediated via intracellular inhibitory G-proteins, which dissociate into *α*- and *βγ*-subunits after formation of a receptor-ligand complex [9]. Among other actions, the *βγ*-subunits bind to calcium channels in the plasma membrane. This leads to closure of the channels, thereby lowering the amount of positive calcium-ion influx and reducing cellular excitability [3, 8, 10, 11].

In this paper, we model this pathway to analyse the effects of fentanyl and NFEPP on the number of closed membrane calcium channels and activated (i.e. dissociated) G-protein complexes at different pH-levels. We constructed a biochemical reaction network that connects the receptor-ligand interactions to the G-protein cycle, and further to the signal cycle of calcium channel opening and closing (see Fig. 1 for an illustration). The corresponding stochastic reaction process was simulated for different values of the receptor-ligand binding rate, comparing the mean inhibition of calcium currents resulting from these numerical simulations to corresponding data from *in vitro* experiments. By numerical simulation, we observe that the binding rate has a non-linear effect onto the mean amplitude of deactivated calcium channels, which explains the different effects of NFEPP and fentanyl in inflamed versus healthy tissue.

**Figure 1:**
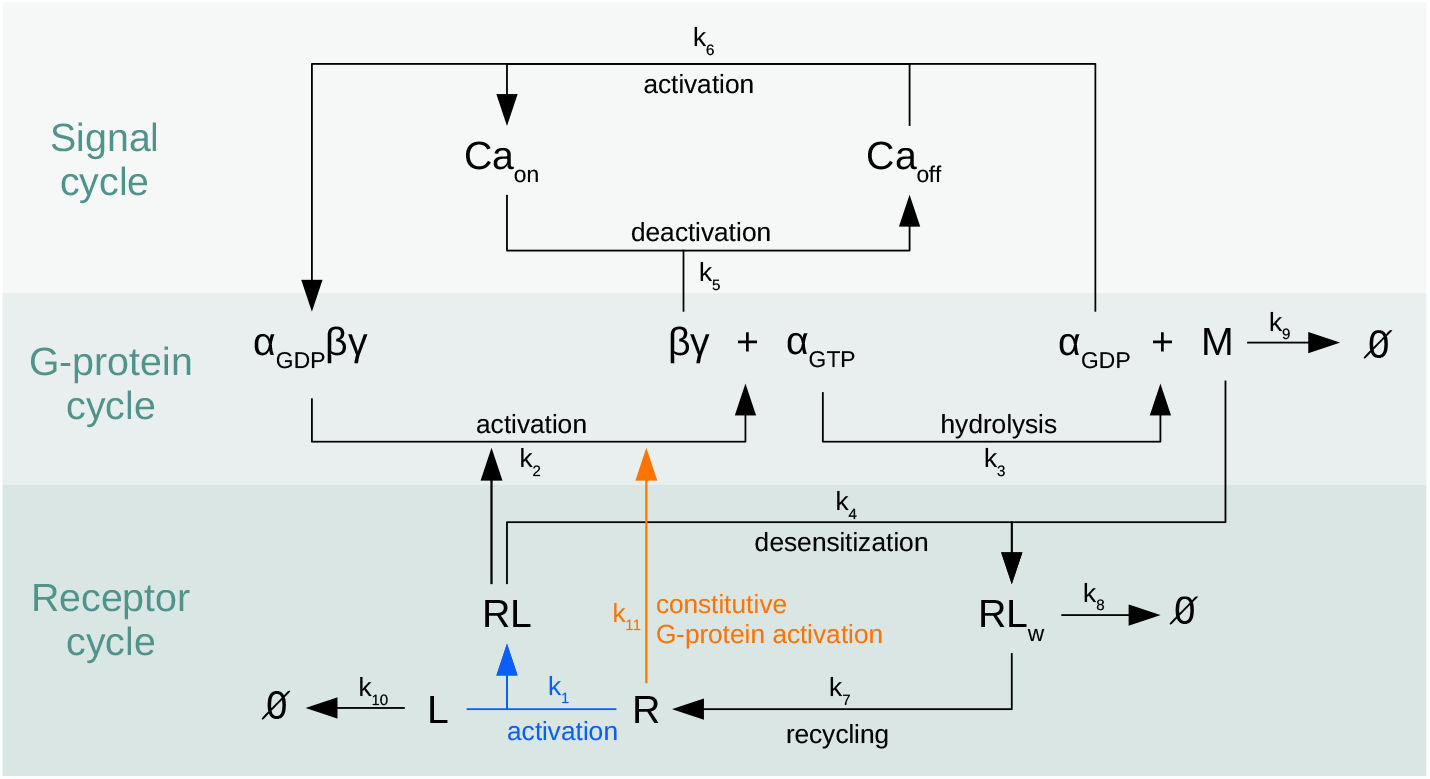
Overview of the reaction network. Biochemical reaction network for the *µ*-opioid receptor signalling pathway, connecting the receptor cycle to the G-protein cycle and further to the signal cycle of membrane calcium channel modulation, see Sec. 2.1 for an explanation. The focal point of this study is the analysis of the impact that the rates for ligand-induced receptor activation (blue) and for constitutive G-protein activation (orange) have onto the overall dynamics.

It is important to note that our approach differs from others that have investigated signalling pathways from receptor to the nucleus or to intracellular second messengers (not to the plasma membrane) [12, 13]. In contrast to those studies, we choose a stochastic approach because it delivers more information than deterministic alternatives.

Aside from pH, other inflammatory mediators play important roles. For example, reactive oxygen species (radicals) can induce disulfide bond (DSB) formation in opioid receptors [14]. In order to understand the interplay between pH and additional DSB inside MOR for the signalling, we modelled different scenarios and performed molecular dynamics (MD) simulations. These simulations showed that DSB inside the binding pocket might initiate G-protein activation in the absence of an opioid ligand (so-called constitutive G-protein activation). Motivated by this observation, we included the reaction of constitutive G-protein dissociation in our network, and performed additional *in vitro* experiments examining G-protein activation by reactive oxygen species.

In summary, two different influences and three different effects are analysed in this article:

1. A lower pH value changes the protonation state of amino acid residues and opioid ligands, and thus may change their binding rates and subsequent modulation of calcium channels (first effect).
2. An increased concentration of radicals leads to DSB formation, which may reduce the binding affinity of ligands (second effect) and may increase the probability for constitutive G-protein dissociation (third effect).

The results suggest that, compared to radicals, low pH is a more important determinant of overall GPCR function in an inflamed environment.

Unlike in our previous work, we here studied these effects on the higher level of molecular interactions: Results from MD and reaction network simulation are integrated with data from *in vitro* experiments in order to analyse the consequences of environment-dependent ligand binding rates onto the downstream signalling, i.e. calcium channel inhibition.

## 2 Models and methods

In this section we introduce a probabilistic model for the signalling pathway from receptor activation over the G-protein cycle to the calcium channel inhibition. We explain the *in vitro* experiments which were performed to validate the modelling results and estimate the parameter values. Moreover, we motivate our choice of parameter values and specify the numerical approach used for solving the system.

### 2.1 The reaction network

The biochemical reaction network under consideration consists of the following reactions (see Tab. 1 for an overview and Fig. 1 for an illustration). A ligand *L* attaches to a receptor *R* in the membrane, resulting in a receptor-ligand complex *RL* (reaction R_1_). This receptor-ligand complex *RL* activates a trimeric G-protein complex which leads to exchange of *GDP* by *GTP* and subsequent dissociation into *α*- and *βγ*-subunits (reaction ℛ_2_). These subunits activate different signalling pathways. Along with the hydrolysis of *GTP*, another reaction partner *M* (e.g. arrestin) emerges (reaction ℛ_3_), which initiates internalisation of the receptor-ligand complex (reaction ℛ_4_). The *βγ*-subunit inhibits a membrane calcium channel by binding to it (reaction ℛ_5_)^1^. After dissociation of the *βγ*-subunit from the calcium channel, a trimeric G-protein complex is reformed, and the calcium channel is opened (reaction ℛ_6_). The internalised receptor *RL*_*w*_ is either recycled to the cell membrane (reaction ℛ_7_) or degraded (reaction ℛ_8_). The reaction partner *M* can itself be degraded (reaction ℛ_9_). The ligand *L* can vanish before it binds to the receptor, e.g. by degradation or unspecific binding to other extracellular components (reaction ℛ_10_), or it is degraded intracellularly (reactions ℛ_7_ and ℛ_8_). The constitutive G-protein activation is given by reaction ℛ_11_, where we simply use ℛ_2_ without ligand.

Given these reactions, the stochastic dynamics of the system are mathematically modelled by a reaction jump process characterised by the chemical master equation [16, 17]. The state of the system is given by a vector

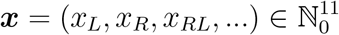

counting the number *x*_*S*_ of molecules of the different species *S ∈* 𝒮, where 𝒮 is the set of species under consideration:

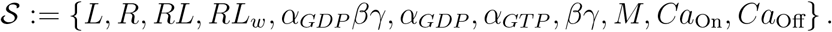

For each reaction ℛ_*j*_ there is a stoichiometric vector **ν**_*j*_ ∈ ℤ^11^ defining the net change in the population state ***x*** induced by this reaction. That is, each time that reaction ℛ_*j*_ occurs, this leads to a jump in the system’s state of the form

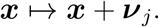

E.g., the stoichiometric vector **ν**_1_ of reaction ℛ_1_ is given by **ν**_1_ = (−1, −1, 1, 0, …, 0). The rates at which the reactions occur are given by propensity functions 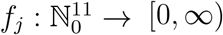, which can be found in the right column of Tab. 1.

The temporal evolution of the system is described by the Markov jump process (***X***(*t*))_*t*≥0_, ***X***(*t*) = (*X*_*S*_(*t*))_*S ∈*𝒮_, where *X*_*S*_(*t*) is the number of molecules of species *S* at time *t*. We define the probability *p*(***x***, *t*) := 𝕡 (***X***(*t*) = ***x*** |***X***(0) = ***x***_0_) to find the system in state ***x*** at time *t* given some initial state ***x***_0_. Then, the overall dynamics are characterised by the standard chemical master equation given by

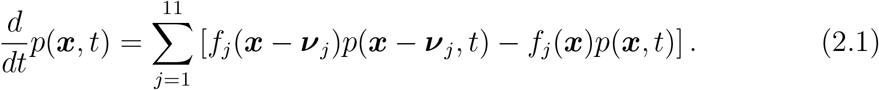

The reaction rate equation characterising the corresponding deterministic reaction system is given by the ordinary differential equation (ODE)

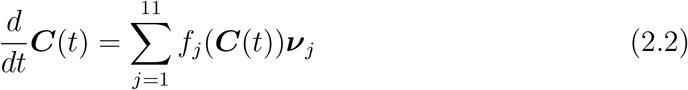

for concentrations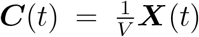, where *V* is the system’s volume. It is well-known that the volume-rescaled Markov jump process (***X***(*t*)*/V*)_0≤*t*≤*T*_ governed by the chemical master equation (2.1) converges to the solution ***C***(*t*)_0≤*t*≤*T*_ of the ODE system (2.2) in the limit of large particle numbers, i.e., for *V* → ∞, see [17] for details.

#### Stochastic vs deterministic approach

The stochastic approach has several advantages over the deterministic one. At first, ODEs are an approximation assuming that the higher moments are trivially given by powers of the first moment. Stochastic modelling is exact in the sense that it takes into account all higher moments. Furthermore, the stochastic approach is closer to reality because it assumes a finite set (discrete number) of molecules, while ODEs consider concentrations and only work as approximations for large particle numbers. So the stochastic model is better suited for modelling a small compartment like an axon terminal with a small number of MORs and G-proteins. Last but not least, a stochastic model delivers more information than ODEs. E.g., it enabled us to analyse the variances of the trajectories or the probability distribution of certain variables like the number of ligand-receptor binding reactions, which will be done in Sec. 3.1.

In many situations, however, the ODE model provides a valid approximation of the rescaled first moment of the stochastic process, ***C***(*t*) ≈𝔼 (***X***(*t*))*/V*, as it is also the case here. This fact will be exploited in Sec. 2.3 where the less complex ODE model instead of the stochastic one will be used for estimating the reaction rates *k*_1_, …, *k*_10_ based on experimental data.

### 2.2 Laboratory *in vitro* experiments

In order to validate our model, we performed laboratory experiments measuring G-protein activation and membrane calcium currents *in vitro*. To determine initial G-protein activation (as reflected by the exchange rate of GDP for GTP), the [^35^S]-GTP*γ*S binding assay was used. Because these experiments require genetic alteration (by transfection) of cells, we performed these measurements in commonly used human embryonic kidney (HEK293) cells. In addition, we extracted data produced by Förster resonance energy transfer (FRET) experiments from [2]. These experiments measure ligand-induced G-protein subunit dissociation (which follows G-protein activation). The FRET experiments were used to fit the reaction rates. To mimic the mechanisms underlying *in vivo* opioid analgesia, we examined calcium currents in sensory neurons harvested from rodents using a patch-clamp protocol (see supporting information for methodological details). The experimental results are shown in Fig. 2 and Fig. 5 below, and described in more detail in Sec. 3.1.

**Figure 2:**
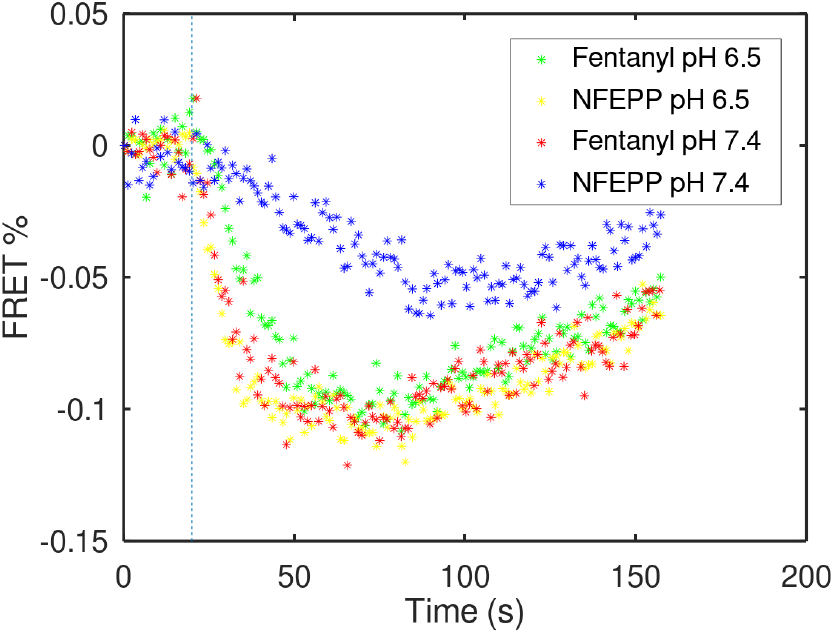
*In vitro* experiments: G-protein activation. Time course of ligand-induced G-protein subunit dissociation measured by FRET in HEK293 cells. FRET efficiency is depicted as percentage of initial intensities, corrected for photobleaching [2]. A higher number of dissociated G-protein subunits (stronger G-protein activation) is represented by more negative values. One can directly see that the blue “curve” (NFEPP at pH 7.4) shows lower numbers of dissociated subunits (weaker G-protein activation) compared to the other scenarios. The dashed line indicates the time point *t* = 20*s* where the ligand was added.

### 2.3 Parameter estimation

Our model includes eleven previously unknown parameters *k*_1_, …, *k*_11_. The determination of appropriate values for these parameters included two main steps: a rough selection of values based on literature, followed by more precise standard parameter estimation.

In [7], it has been shown by means of MD simulations that the ligand binding affinity varies for different ligands and pH values. Based on these results, we chose the following different values of *k*_1_ for different ligand/pH combinations: *k*_1_ = 1.25 × 10^−2^*s*^−1^ for fentanyl and pH 6.5, *k*_1_ = 2.5 × 10^−2^*s*^−1^ for fentanyl/pH 7.4, *k*_1_ = 2.5 × 10^−2^*s*^−1^ for NFEPP/pH 6.5, and *k*_1_ = 5 × 10^−4^*s*^−1^ for NFEPP/pH 7.4. As the *in vitro* experiments used for the parameter estimation were performed in the absence of radicals, the rate *k*_11_, which is responsible for the constitutive G-protein activation ℛ_11_ was set to zero.

The parameters *k*_2_, …, *k*_10_ of the other intracellular reactions were assumed to depend only mildly (if at all) on the ligand/pH combination in the cellular environment; for each of these parameters a single value has been chosen, independent of ligand and pH. This is a reasonable assumption because we chose an intracellular pH value of 7.4 based on well-known mechanisms of cellular homeostasis: Although transient (several minutes) changes of intracellular pH may occur with tissue acidosis, intracellular buffer systems and ion pumps in the plasma membrane will rapidly restore physiological pH to ensure cell viability [18]. Since most previous studies examined situations of longer-lasting inflammation (up to several days) [2, 4, 5, 6], we look at this situation, as well. Using results from [19], we started by setting the rate constant *k*_5_ of the central binding reaction between the βγ-subunit and the calcium channel to *k*_5_ = 5 × 10^−2^*s*^−1^ and proceeded to arrange the other values relative to it according to what is known in the literature. Comparing [19] and [20] it can be deduced that ℛ_5_ happens at a timescale an order of magnitude shorter than ℛ_2_ and ℛ_3_ (with ℛ_2_ being slightly faster than ℛ_3_), so we chose *k*_2_ and *k*_3_ five resp. ten times smaller than *k*_5_. The rate constants of reactions ℛ_4_, ℛ_6_, ℛ_9_ were assumed to be of the same magnitude as those of ℛ_2_ and ℛ_3_ (with ℛ_6_ being slightly faster). The recycling and degradation of internalised ligand-receptor complexes are much slower, at a level of minutes (Fig. 1 in [21]) which leads to comparatively small rate constants *k*_7_ and *k*_8_ for the reactions ℛ_7_ and ℛ_8_ of the internalised receptor. The extracellular decay of ligand due to unspecific binding and other incidents (reaction ℛ_10_) was set to a value at which it showed a first effect on calcium channel inhibition.

After this initial step of selecting rough parameter values based on available information, we fine-tuned the parameters *k*_1_, …, *k*_10_ via standard parameter estimation techniques using the in-vitro experimental FRET data (see Fig. 2 below) that consists of four individual time series for G-protein activation for the four cases fentanyl/pH 6.5, fentanyl/pH 7.4, NFEPP/pH 6.5, and NFEPP/pH 7.4. The experimental data was first pre-processed by determining the offset time (time point at which the respective ligand was added), and the linear scaling transformation that maps the number of undissociated G-proteins from the model to the measured FRET signal. Then the residual distance between the solution of the ODE model and the experimental data was minimized by optimally adapting the parameters *k*_2_, …, *k*_10_, starting from the initial values previously chosen (second column of Table 2 termed “Preselected”). Here, the residual is the mean squared distance between ODE solution and data, summed over all four time series. The minimization was done using standard techniques for parameter estimation [22, 23], within the framework of the software PREDICI [24]. The resulting parameters values are shown in Table 2, third column termed “Parameter Estimation”. These values of *k*_2_, …, *k*_10_, optimally adapted to all four cases of ligand/pH combination at once, were fixed.

**Table 1:** Reactions and propensity functions. For each reaction ℛ_*j*_ there is a propensity function *f*_*j*_ giving the rate (probability per unit of time) for the reaction to occur depending on the system’s state ***x*** = (*x*_*L*_, *x*_*R*_, …). For any species *S* it stands *x*_*S*_ for the number of molecules of this species. *R*: receptor, *L*: ligand, *RL*: receptor-ligand complex, *RL*_*w*_: internalised receptor, *α*_*GDP*_ *βγ*: G-protein, *α*_*GDP*_ */α*_*GT P*_: *α*-subunit loaded with *GTP* or *GDP*, respectively, *βγ*: *βγ*-subunit, *M*: reaction partner (e.g. arrestin) to initiate receptor internalisation, *Ca*_Off_*/Ca*_On_: closed/open calcium channel.

| $j$ | Reaction $\mathcal{R}_j$ | Propensity $f_j$ |
| --- | --- | --- |
| 1 | $L + R \xrightarrow{k_1} RL$ | $k_1 \cdot x_R \cdot x_L$ |
| 2 | $RL + \alpha_{GDP}\beta\gamma \xrightarrow{k_2} RL + \alpha_{GTP} + \beta\gamma$ | $k_2 \cdot x_{RL} \cdot x_{\alpha_{GDP}\beta\gamma}$ |
| 3 | $\alpha_{GTP} \xrightarrow{k_3} \alpha_{GDP} + M$ | $k_3 \cdot x_{\alpha_{GTP}}$ |
| 4 | $RL + M \xrightarrow{k_4} RL_w$ | $k_4 \cdot x_{RL} \cdot x_M$ |
| 5 | $\beta\gamma + Ca_{On} \xrightarrow{k_5} Ca_{Off}$ | $k_5 \cdot x_{\beta\gamma} \cdot x_{Ca_{On}}$ |
| 6 | $\alpha_{GDP} + Ca_{Off} \xrightarrow{k_6} \alpha_{GDP}\beta\gamma + Ca_{On}$ | $k_6 \cdot x_{\alpha_{GDP}} \cdot x_{Ca_{Off}}$ |
| 7 | $RL_w \xrightarrow{k_7} R$ | $k_7 \cdot x_{RL_w}$ |
| 8 | $RL_w \xrightarrow{k_8} \emptyset$ | $k_8 \cdot x_{RL_w}$ |
| 9 | $M \xrightarrow{k_9} \emptyset$ | $k_9 \cdot x_M$ |
| 10 | $L \xrightarrow{k_{10}} \emptyset$ | $k_{10} \cdot x_L$ |
| 11 | $R + \alpha_{GDP}\beta\gamma \xrightarrow{k_{11}} R + \alpha_{GTP} + \beta\gamma$ | $k_{11} \cdot x_R \cdot x_{\alpha_{GDP}\beta\gamma}$ |

**Table 2:**
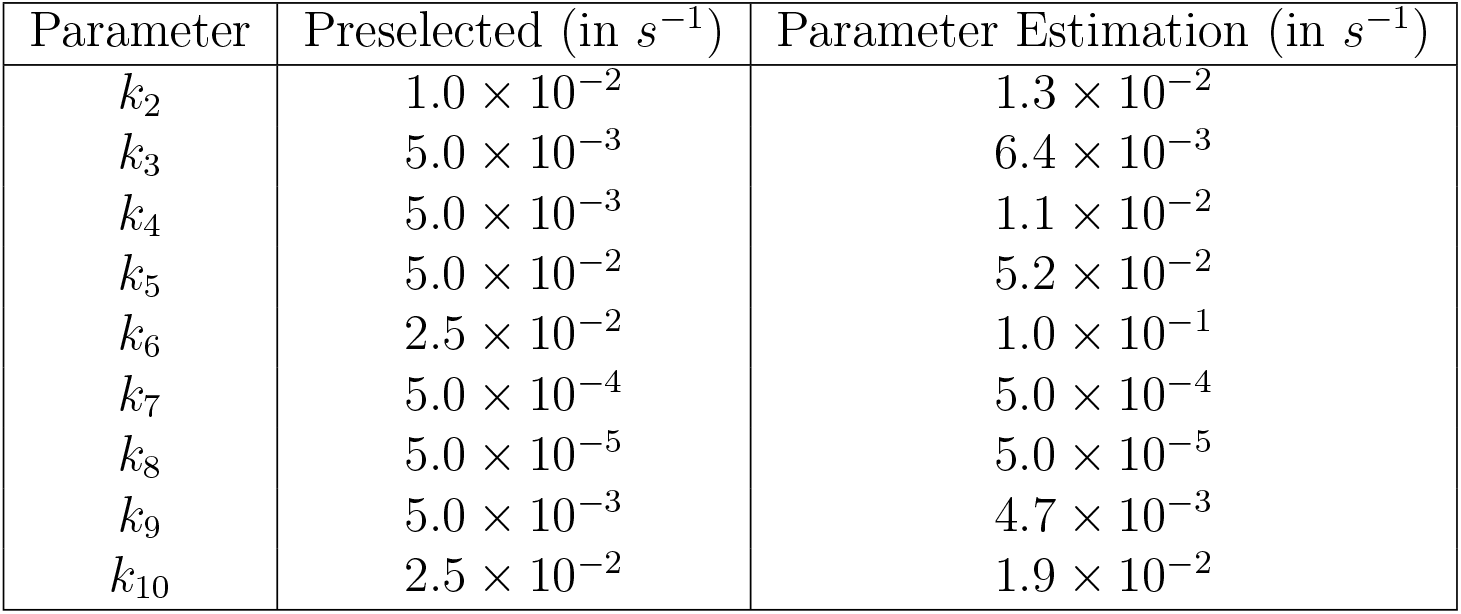
Reaction rate constants. Optimal parameter values compared to the assumed values. The parameter *k*_11_ is not listed here because its value is assumed to be fixed to *k*_11_ = 0. For *k*_1_ see text.

In a final step, individually for each ligand/pH combination, the parameter *k*_1_ was fine-tuned by minimizing the residual function for each single time-series for fixed *k*_2_, …, *k*_10_ by changing the respective *k*_1_. The outcome was:

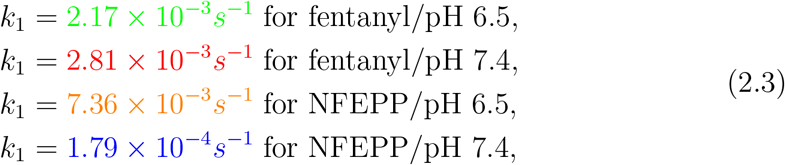

where colors shall help to identify the respective curves in Figs. 3-5.

**Figure 3:**
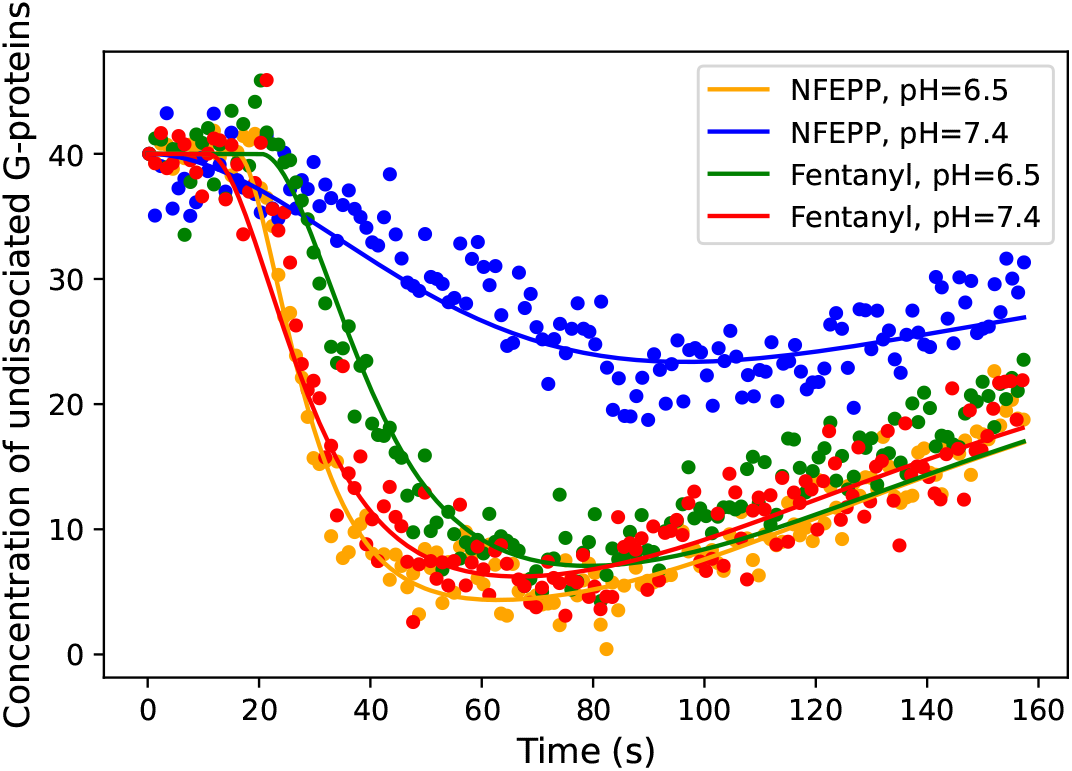
*In vitro* data and optimally fitted ODE model. Dots represent the time course of ligand-induced G-protein subunit dissociation measured by FRET in HEK293 cells. FRET values were transformed into concentration of undissociated G-proteins by a scaling factor. Lines indicate the best-fit of the ODE model to the data (using optimal parameters). For methodological details, see [2].

Some of the optimal parameter values exhibit mild deviations from the pre-selected ones, but no stark contrast to the literature was observed; in fact, closer inspection showed that the mean squared deviation between model-based simulation and experiment was significantly reduced by fine-tuning parameters. The resulting best fit is shown in Fig. 3.

It has been tested that, for these parameter values, the residual distance between data and stochastic model is very close to the residual distance for the ODE model.

### 2.4 Numerical simulations

Simulations of the stochastic reaction kinetics were made with Python 3. For each combination of rate constants, 500 Monte Carlo simulations were carried out and the arithmetic mean was calculated in order to estimate the percentage of closed calcium channels plotted in Fig. 4 and Fig. 12. As a time horizon for each simulation 1200 seconds were chosen. The results are presented in Sec. 3. In order to find the steady state of the dynamics for non-zero *k*_11_, a simulation without ligands (or, equivalently, with *k*_1_ = 0) was run. This long-term mean *a* (rounded to natural numbers) of closed calcium channels was then used to determine the initial state for the dynamics including receptor activation (see Tab. 3 for the results). To check for normal distribution of the mean, the 500 runs were divided into batches of 50 and the respective means then tested. Anderson-Darling test indicated normal distribution with *p* ≤ 0.05, so the 95%-confidence interval of the t-distribution is shown in the plots.

**Figure 4:**
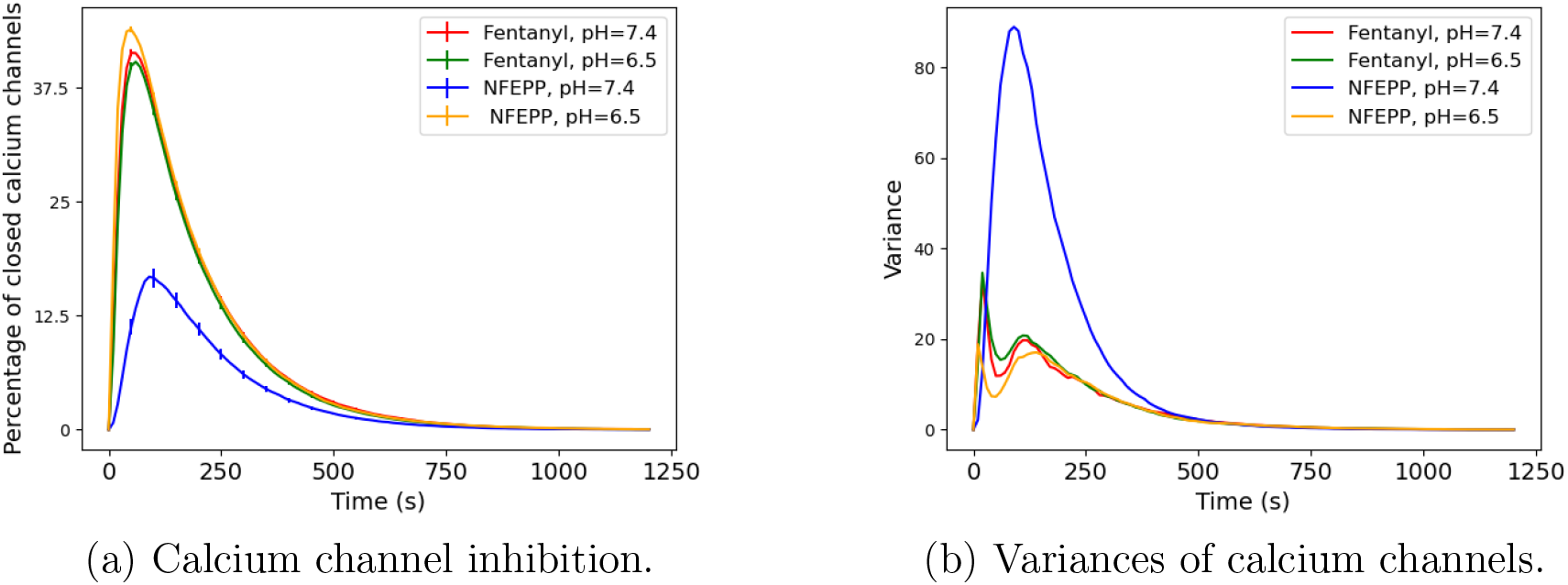
Numerical studies. (a) Time course of percentage of closed calcium channels for different values of the ligand-binding rate *k*_1_. Error bars indicate 95%-confidence intervals (from 500 simulation runs). (b) Variances of closed calcium channels from 500 simulation runs.

## 3 Results

The reaction network introduced in Sec. 2 was used to analyse the impact (a) of different extracellular pH values correlating to those occurring in injured tissues *in vivo*[3] in combination with a conventional or a pH-dependent opioid ligand (fentanyl or NFEPP, respectively) (see 3.1), and (b) of oxygen radicals and additional DSB (see 3.2) onto the overall signalling pathway. Based on our parameter fitting results, the changing pH value was modelled via varying the rate constant *k*_1_ of ligand-receptor binding. As for the impact of additional DSB, MD simulations showed an effect on both *k*_1_ and on the rate constant *k*_11_ for constitutive G-protein activation.

### 3.1 Isolated impact of pH value

For the following analysis we set the rate constant *k*_11_ to zero. Our goal was to analyse the effect of varying rates *k*_1_ *>* 0 for the ligand-induced activation of a receptor (given by the binding reaction ℛ_1_: *L* + *R* → *RL*) onto the amount of closed calcium channels *Ca*_Off_. We examined the ligands fentanyl and NFEPP in combination with changing pH levels (see equation (2.3) for the respective rate values *k*_1_). The rates for the other reactions were left unchanged in all simulations based on the assumption that intracellular pH remains at 7.4 (see Table 2).

Fig. 4a represents the evolution of the mean number of closed calcium channels for the different ligand-binding rates *k*_1_ given in equation (2.3). For all ligand and pH pairs except for NFEPP/ pH=7.4 we observed similar amplitudes of closed calcium channels (about 44% of all calcium channels), while for NFEPP/ pH=7.4 the amplitude is significantly reduced to approximately 29% (but note that only a maximum of 40 out of the total of 80 channels can be closed since there are only 40 G-proteins, so the maximum calcium channel inhibition is 50%). In Fig. 4b the variances of the number of closed calcium channels depending on time is shown.

We can observe that this variance is significantly larger for NFEPP at a normal pH value than for all other scenarios.

#### Results from *in vitro* experiments

The maximum inhibition of voltage-induced calcium currents by fentanyl or NFEPP at pH 6.5 and pH 7.4 was measured by patch-clamp experiments in rat sensory neurons. The results are comparable to the scenarios simulated in Fig. 4a in that both fentanyl and NFEPP potently inhibited calcium currents at low pH, whereas NFEPP was significantly less effective than fentanyl at normal pH (Fig. 5). Fig. 2 shows the experiment that was used for data fitting.

**Figure 5:**
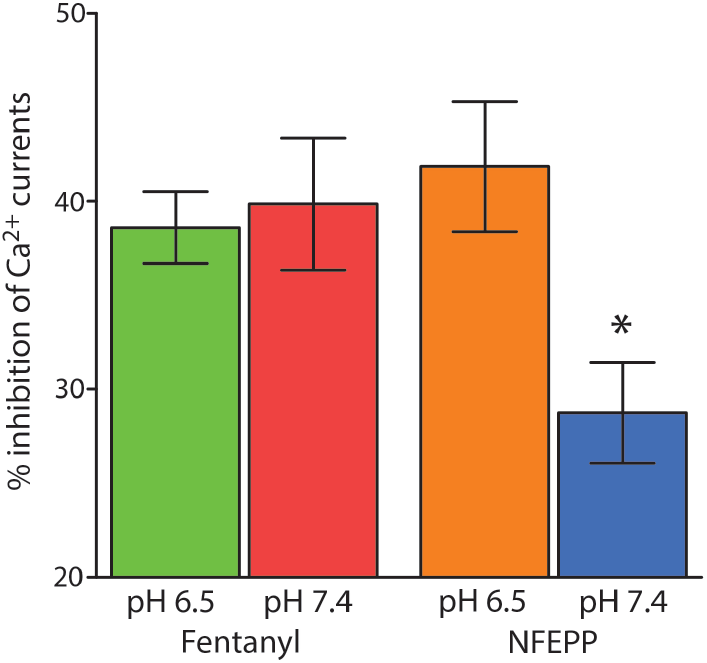
*In vitro* experiments: Calcium currents. Maximum inhibition of voltage-induced calcium currents by fentanyl or NFEPP at pH 6.5 and pH 7.4 measured by patch clamp experiments in rat sensory neurons. *P < 0.05 for comparison of NFEPP at pH 7.4 to all other values (one-way ANOVA with Bonferroni’s multiple comparisons). Data are means ± standard error of the mean (SEM).

#### Non-linear behaviour and stochastic effects

Seeing that our model resembled the *in vitro* results quite well, we now sought to get more information about the dependence of the calcium channel inhibition amplitude and the binding rate *k*_1_. Therefore we plotted both against each other; the results are shown in Fig. 6. We see a non-linear behaviour of the calcium channel inhibition amplitude with a rather sharp drop for *k*_1_ < 2×10^−3^*s*^−1^ where the calcium channel response declines quickly.

**Figure 6:**
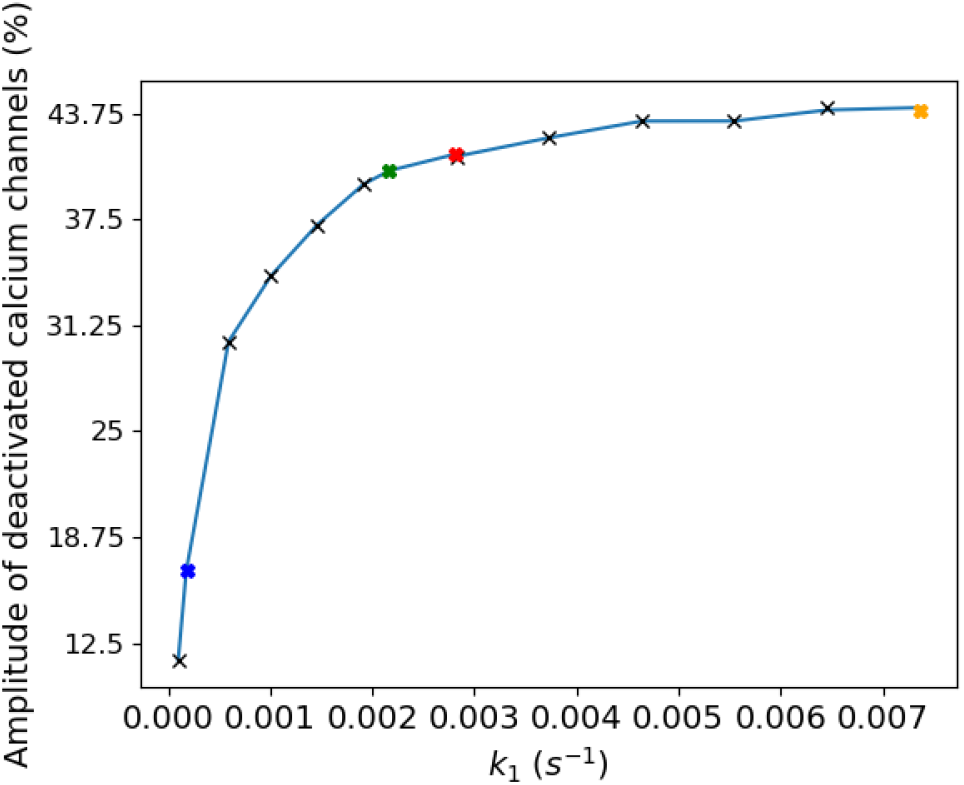
Numerical studies. Amplitude of the mean calcium channel inhibition plotted against *k*_1_. Coloured crosses indicate the *k*_1_ values according to equation (2.3).

In order to investigate the stochastic effects, we calculated the probability distributions of the number of binding reactions that happened during a simulation run over the time interval [0, 1200*s*] (see Fig. 7). For decreasing *k*_1_ (see 2.3 for the corresponding ligand/ pH pairs) the distribution shifts to lower values and gets a wider range. The non-linearity is also represented in the larger shift from (c) to (d) compared to the other shifts.

**Figure 7:**
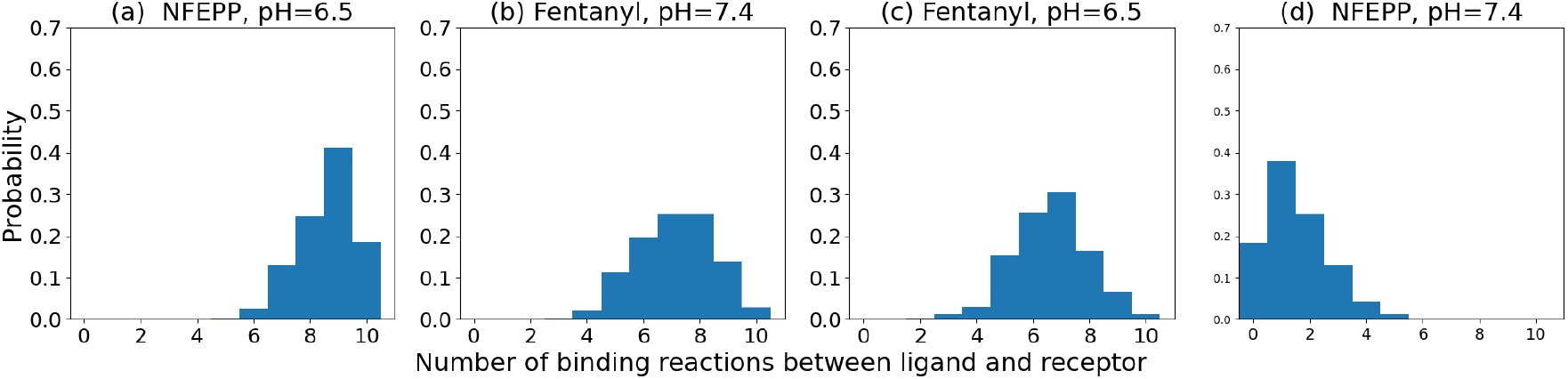
Probability distribution of the number of receptor-ligand binding reactions ℛ_1_ that happened during the whole time course of a simulation run (time interval [0, 1200*s*]). For each *k*_1_-level, 500 simulation runs have been evaluated.

### 3.2 Combined impact of pH value and additional DSB

We next investigated the effect of rising radical levels (resulting in additional DSB in the receptor) in combination with varying pH values onto the dynamics. The results of the MD simulations are described in Sec. 3.2.1, while the consequences for the reaction process will be presented in Sec. 3.2.2.

#### 3.2.1 Results from MD simulations

We saw that the pH value has an impact on the binding rate of NFEPP and fentanyl. However, we found that the receptor-ligand binding rate was also influenced by an additional DSB, which is typically promoted by increased radical concentrations [14]. In the atomistic MD simulations, the difference between inflamed and healthy tissue was modelled by changes in pH and with the formation of a DSB. In order to account for protonation and deprotonation of respective amino acid residues and ligands, the simulation parameter setting for inflamed tissue was pH 5 and the setting for healthy tissue was pH 7. As explained in the supporting information (Sec. A.1), this parameter setting does not really represent a concrete *H*^+^-ion concentration, but only has an influence on the protonation state of the MOR amino acid residues. The setting pH 7 results in the same protonation states of amino acid residues as pH 7.4 and, therefore, models the healthy tissue situation. In the rat MOR, CYS 292^6.47^ of the TM6 helix along with CYS 321^7.38^ of the neighbouring TM7 helix were selected for the introduction of an additional DSB. Sulfur atoms of these two cysteine residues are at a distance of 0.987 nm in the native rat MOR crystal structure according to Protein Data Bank (PDB) [25], code 6DDF [26]. Significantly, CYS 292^6.47^ is also in proximity of HIS 297^6.52^, which is crucial for the interaction of the binding pocket of the receptor with opioid ligands [7, 26]. Hence, it is of special interest to examine ligand binding and activation of the MOR without and with the additional CYS 292^6.47^ – CYS 321^7.38^ DSB in the receptor, as depicted in Fig. 8.

**Figure 8:**
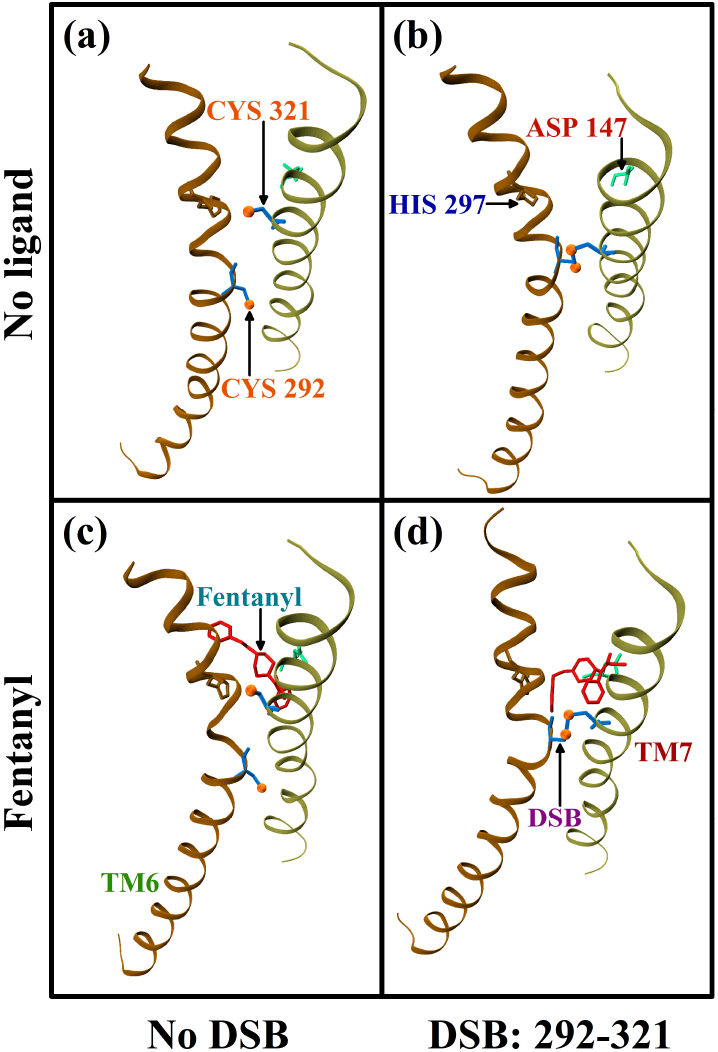
Simulation setup. Trajectory snapshot at 5 ns of *µ*-opioid receptor (MOR) without (a, b) and with (c, d) the ligand fentanyl, and without (a, c) and with (b, d) an additional CYS 292^6.47^ – CYS 321^7.38^ DSB (for terminology, see [27]). The figures show the two neighbouring helices TM6 and TM7 of the MOR. Interaction between the ligand and the MOR is assessed by measuring the distance between the ligand, and the two amino acid residues ASP 147^3.32^ and HIS 297^6.52^.

##### Receptor-ligand interaction

The binding of an opioid ligand to the MOR occurs mainly between the two amino acids ASP 147^3.32^ and HIS 297^6.52^ [26, 27]. The ligand positions itself between these two residues. Thus, the interaction of the binding region of the MOR with the ligand was assessed by measuring its distance with regard to the crucial ASP 147^3.32^ and HIS 297^6.52^ residues of the binding region [7].

The formation of an additional DSB is promoted by radicals and, thus, due to the situation of inflamed tissue. In healthy tissue the formation of additional DSB is unlikely. This means, that in Fig. 9 mainly 9a and 9c are of importance. At pH 5, fentanyl exhibited similar interactions with the binding region irrespective of the additional DSB, as shown in Fig. 9a. However, the fluctuation in the receptor-ligand interaction was demonstrably higher without the extra DSB. The ligand stays in greater proximity of ASP 147^3.32^ as compared to HIS 297^6.52^. For NFEPP, interaction with the ASP 147^3.32^ residue at pH 5 gets affected upon the introduction of the additional DSB. However, ligand interaction with HIS 297^6.52^ remains similar for both scenarios (Fig. 9c). From this observation, we conclude that the presence of an additional CYS 292^6.47^ – CYS 321^7.38^ DSB has an effect on the binding mode of opioids. An additional DSB can have a significant influence on these systems especially in the case of NFEPP. Hence, increased concentrations of radicals (which induce the formation of DSB) can indeed affect ligand binding at the MOR and perhaps the subsequent signalling events downstream. Our conclusion from the change in the atomic distances between the opioid ligands and the important binding positions is that DSB formation reduces the binding rate *k*_1_. A similar role of DSB has been previously implicated in the modification of the ligand-access channel of cytochrome P450 2B1 [28] and in the functionality of other GPCRs [9].

**Figure 9:**
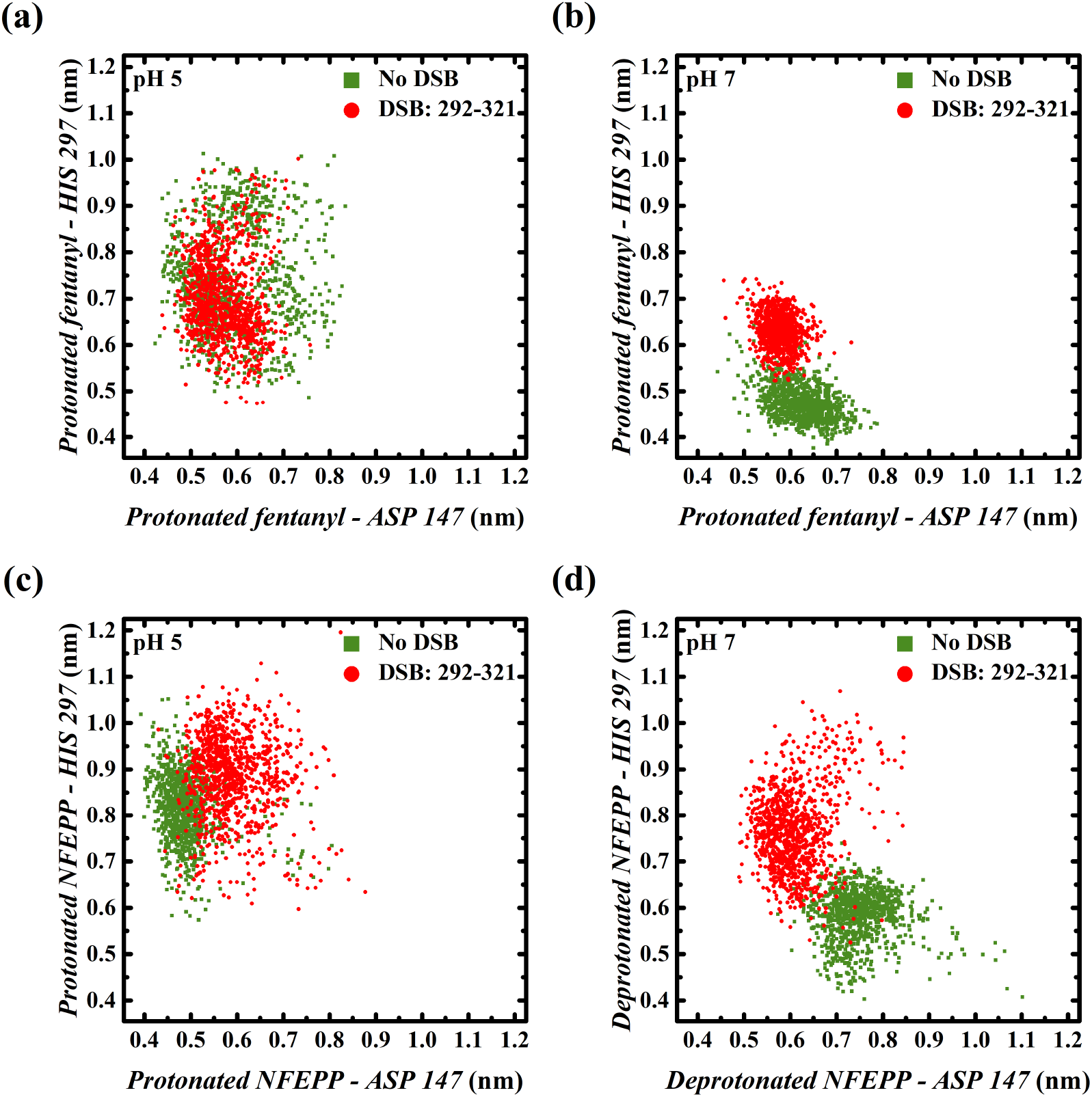
Receptor-ligand interaction. Distance distributions of fentanyl (a, b) and NFEPP (c, d) as a function of system acidity, with respect to the ASP 147^3.32^ and HIS 297^6.52^ residues of the binding region. System states without and with an additional CYS 292^6.47^ – CYS 321^7.38^ DSB are represented by green filled-squares and red filled-circles, respectively.

In inflamed tissue, the amino acid residues are not “completely protonated”. With a low probability, we also find the situations that correspond to a parameter setting of pH 7 in molecular simulations. Fentanyl would interact less with HIS 297^6.52^ with an additional DSB in the receptor using this parameter setting. However, the distance of the ligand from ASP 147^3.32^ remains similar, both without and with the extra DSB (Fig. 9b). NFEPP prefers interaction with HIS 297^6.52^ without the DSB, and with ASP 147^3.32^ if an extra DSB would be present at pH 7 (Fig. 9d). This again shows that a lower rate constant *k*_1_ can be expected in the case of DSB formation.

##### Constitutive G-protein dissociation

The TM6 of the MOR is known to play a crucial role in ligand binding. Furthermore, the outward movement of TM6 is the largest structural change upon receptor activation [9]. The position of TM6 may change just because of the presence of an additional DSB, even if a ligand is not bound. Changes in the MOR conformation were monitored by tracking the distance between the TM6 and TM7 helices in MD simulations without a ligand, as depicted in Fig. 10. The additional CYS 292^6.47^ – CYS 321^7.38^ DSB causes a reduction in the distances between these two helices by approximately 0.1 nm at both pH 5 and 7. Hence, a conformational change of MOR might occur in inflamed tissue, as the surrounding environment turns more acidic accompanied by increased radical concentrations, which can trigger formation of DSB [14].

**Figure 10:**
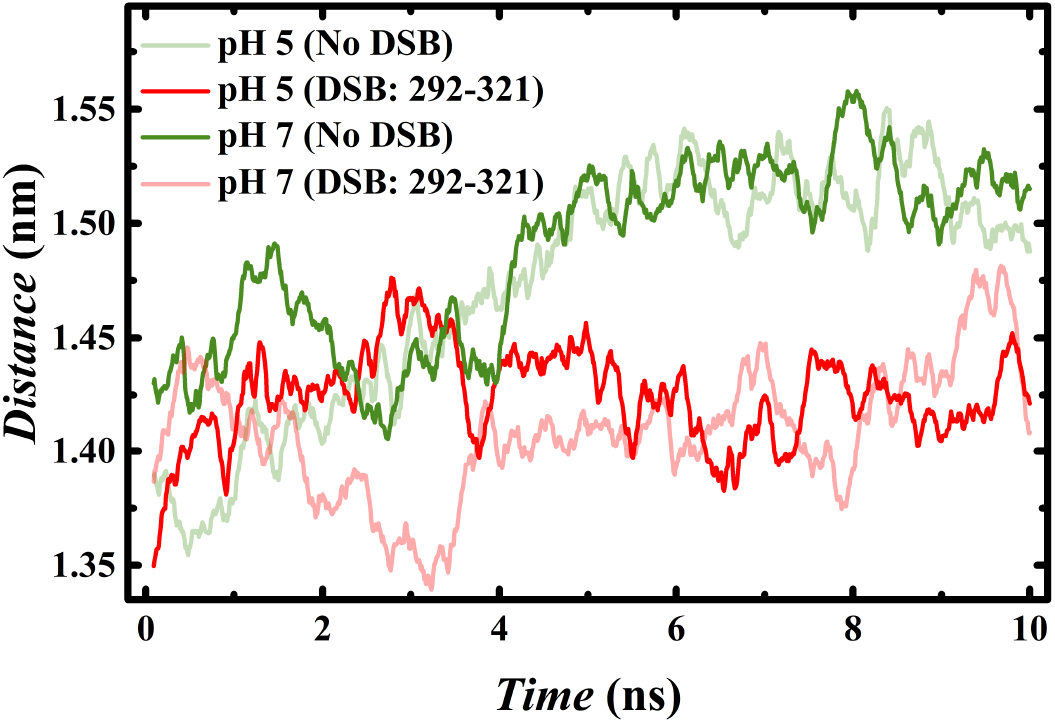
MOR conformation. Time evolution of distance between TM6 and TM7 helices as a function of system pH values of 5 and 7, and absence (green) or presence of an additional CYS 292^6.47^ – CYS 321^7.38^ DSB (red). Physiologically relevant and relatively transient states are represented by solid and faded lines, respectively.

From our MD simulations we see that the position of TM6 depends on several extracellular factors. It is conceivable that DSB-induced conformational changes of TM6 influence the spontaneous (constitutive) dissociation of G-protein subunits in the absence of an opioid ligand. So our MD simulations imply that we can include the possible influence of DSB by changing the rate *k*_11_ for constitutive activation.

##### Results from *in vitro* experiments

The formation of DSB is chemically based on reactive oxygen species [14]. These species can be produced in *in vitro* experiments by adding H_2_O_2_ to the sample (see supporting information for methodological details). Our experimental data support that increasing radical (H_2_O_2_) concentrations (likely associated with increasing DSB formation) are correlated with increasing constitutive G-protein activation (Fig. 11).

**Figure 11:**
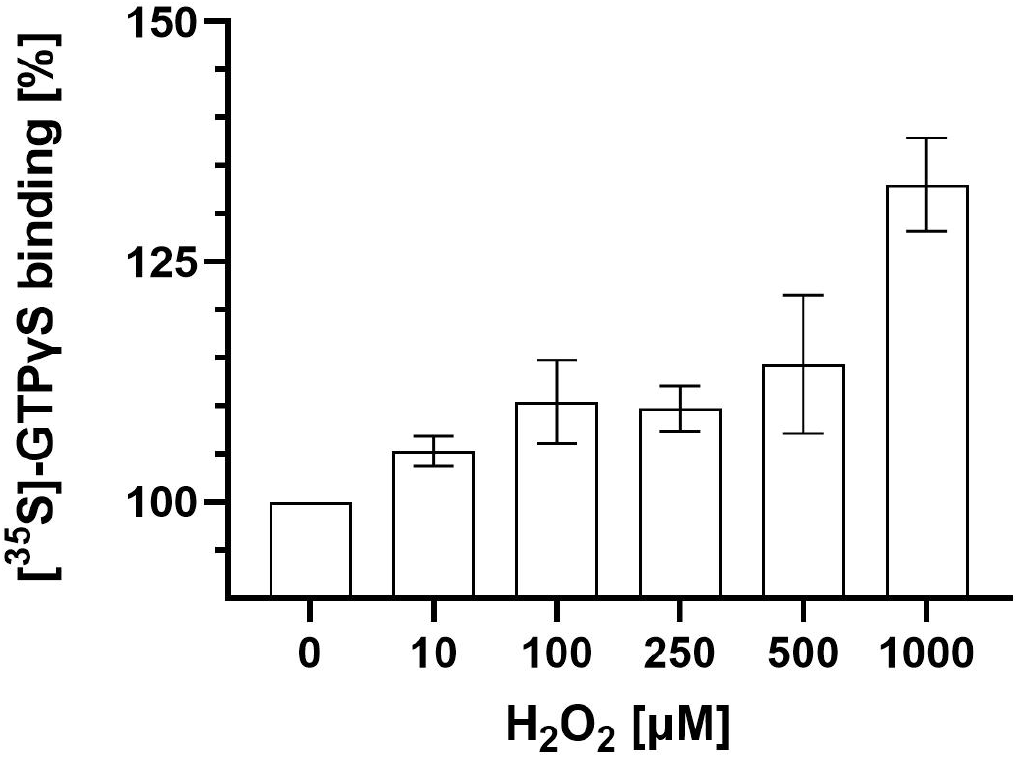
*In vitro* results on constitutive G-protein activation. Effects of increasing concentrations of radicals (H_2_O_2_) on basal [^35^S]-GTP*γ*S binding to MOR without opioid ligands. Data are means ± SEM of specific binding normalised to the control group; *n* = 8 per condition. *P* < 0.05, linear regression analysis.

#### 3.2.2 Effects onto the signalling pathway

The results from the MD simulations described before showed that oxygen radicals (inducing additional DSB) lead to a decrease of the receptor-ligand binding rate *k*_1_ and an increase in constitutive G-protein activation modelled with rate *k*_11_. With progressive inflammation, there are now two effects on *k*_1_, one from the pH and one from DSB. Based on the MD results we estimated that the additional DSB effect decreases the values for *k*_1_ given in equation (2.3) to 80% at pH 6.5. For the concrete values see Table 4.

**Table 3:** Initial state. Initial number *X*_*S*_(0) of molecules for each species *S* ∈ 𝒮 used for all simulations, where *a* = 0 for *k*_11_ = 0 (Sec. 3.1) and *a* = 5 for *k*_11_ = 5 × 10^−5^ (Sec. 3.2)

| Species $S$ | $L$ | $R$ | $RL$ | $RL_w$ | $\alpha\beta\gamma$ | $\alpha_{GDP}$ | $\alpha_{GTP}$ | $\beta\gamma$ | $M$ | $Ca_{On}$ | $Ca_{Off}$ |
| --- | --- | --- | --- | --- | --- | --- | --- | --- | --- | --- | --- |
| $X_S(0)$ | 10 | 20 | 0 | 0 | $40 - a$ | $a$ | 0 | 0 | 0 | $80 - a$ | $a$ |

**Table 4:**
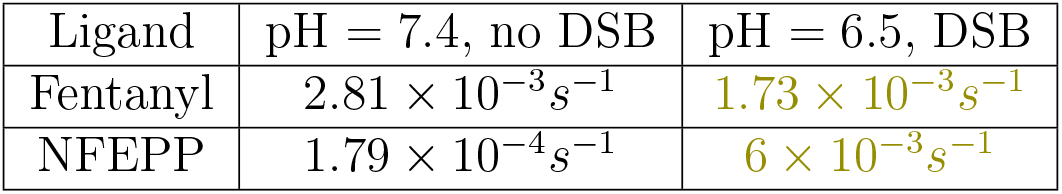
Receptor-ligand binding rates. Rate constant *k*_1_ for receptor activation by ligand-binding, depending on the ligand, the pH-level and DSB presence. These values result from the ones given in equation (2.3) after multiplying by the factor 0.8 for the inflamed scenario (pH = 6.5, additional DSB) while leaving unchanged for pH = 7.4 and no DSB.

The rate *k*_11_ for constitutive G-protein activation was set to *k*_11_ = 5 × 10^−5^ in the presence of DSB. This induces a base level of approximately 5 closed calcium channels in the healthy tissue scenario, which appears to be a reasonable value compared to the other model parameter values. As there was no *in vitro* experimental data available for the scenario including oxygen radicals, a parameter estimation for *k*_11_ was not possible. In case of no DSB we set *k*_11_ = 0 as before. According to the resulting model, the base level of closed calcium channels increases with progressive inflammation (i.e. rising radical concentrations). This is seen in Fig. 12 which shows two plots, one for fentanyl and one for NFEPP. The black curve represents the healthy tissue situation (pH 7.4, no DSB) while the olive (pH 6.5, DSB) curve shows the effects of progressive inflammation. The change of *k*_1_ has the already known effect of reducing the amplitude of the closed calcium channels which is not altered by the constitutive receptor activity.

**Figure 12:**
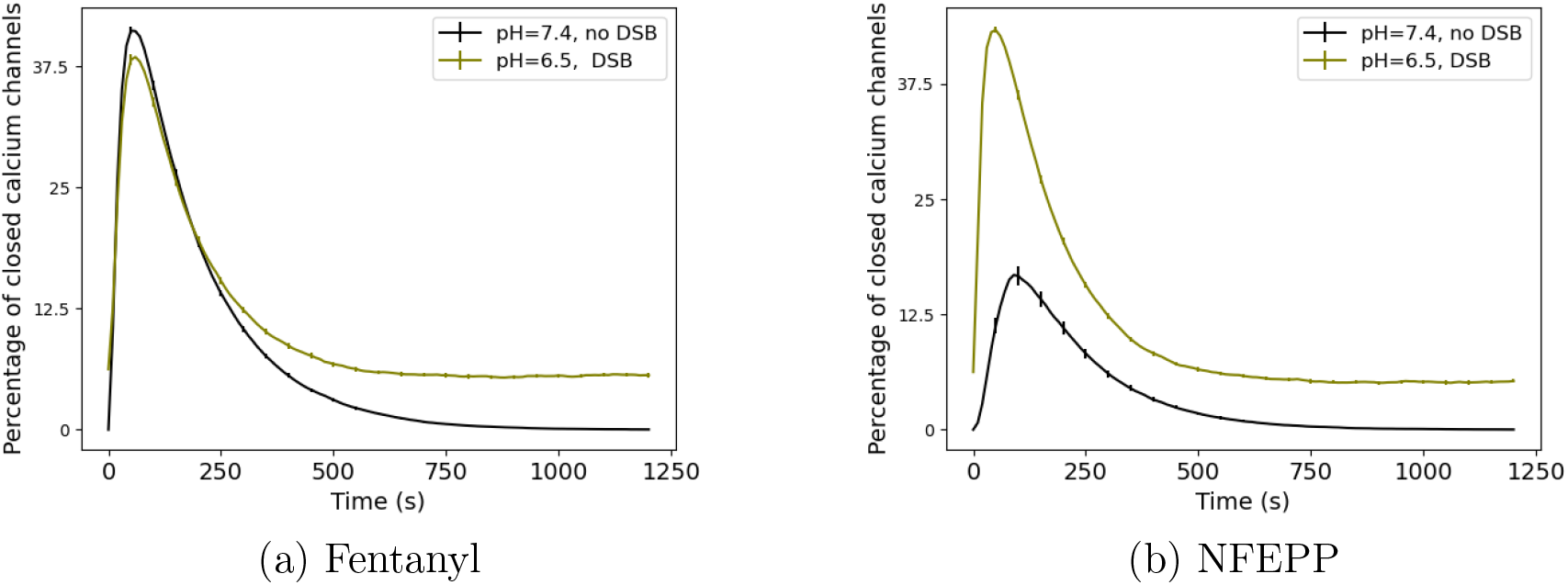
Effects of ligand-binding rate. *k*_1_ **and constitutive activation rate** *k*_11_. Time course of percentage of closed calcium channels for different *k*_1_- and *k*_11_-values for the ligands fentanyl and NFEPP (*k*_11_ = 0 for the case of no DSB and *k*_11_ = 5 × 10^−5^ in case of DSB presence, while the values of *k*_1_ are given in Table 4). Black curves represent the healthy tissue situation, while olive shows the effects of more inflammation (lower pH, more receptors with DSB). The time axis is the same as in Fig. 4a. Error bars indicate 95%-confidence interval (for 500 simulation runs).

## 4 Discussion

We presented a stochastic model of a canonical GPCR signalling pathway linked to plasma membrane function. This pathway is composed of a biochemical reaction network which begins at the receptor, continues with the G-protein and extends to the membrane calcium channels. The respective reaction rates were determined by fitting the model to data from *in vitro* experiments. In addition, we have studied the functional role of DSB inside the receptor’s binding pocket. Our modelling results regarding calcium channels were validated by an independent *in vitro* experiment.

We showed that the change in reaction rates translates into a markedly diminished effect (i.e. a lower number of closed calcium channels) of NFEPP at normal pH compared to all other scenarios (NFEPP at low pH, fentanyl at low or normal pH). The validated model shows a non-linear behaviour of the calcium channel inhibition response with regard to the change of the receptor-ligand binding rate *k*_1_. A critical value of *k*_1_ at which the response drops markedly, is *k*_1_ = 2 × 10^−3^*s*^−1^ (see Fig. 6). In the vicinity of this value, small changes of *k*_1_ lead to large changes of calcium channel inhibition response. So when the pH is sinking below a certain level, the dependence on pH for NFEPP results in a quickly decreasing effect of NFEPP. These results support our previous studies demonstrating that the conventional ligand fentanyl activates MOR both in injured (low pH) and non-injured (normal pH) tissues, while NFEPP is not active in non-injured environments (brain, intestinal wall) [2, 4, 5, 6].

We were able to investigate this phenomenon further and can state that this decrease is not due to a uniform decrease of all stochastic realisations of the reaction process but to a stronger decrease of some realisations and the nearly unchanged course of others. Mathematically, this is represented by the rise of the variance (see Fig. 4b).

These results are an extension of the findings in our earlier work [2]. There it was theorised and corroborated in animal studies that a ligand with proper pH-dependent binding rate would exhibit analgesic effects without side effects. Now we can add that the change of binding rates results in reduced calcium channel inhibition. Thus, the present data provide a more detailed explanation by including the intracellular signalling pathway underlying our initial findings. This further supports our concept of targeting disease-specific conformations of MOR to preclude adverse side effects of painkillers.

To find out about the effect of other inflammatory mediators, in our case additional DSB, a two stage approach was used: As a first step, qualitative changes of the reaction rates were assessed by MD simulations, and as a second step, these changes were used for stochastic simulations of the corresponding reaction process. The MD simulations with DSB imply a decrease of *k*_1_. With the amount of decrease observed, no decisive changes in the effect of both ligands are to be expected. Only if one assumes the decrease to be large enough to drop *k*_1_ below the critical value of 2 × 10^−3^*s*^−1^, marked changes will be seen. This was also observed based on the reaction process model. Moreover, the MD simulations give evidence for a relatively higher constitutive receptor activity which could also be seen in *in vitro* experiments. Absolute values cannot be inferred from the current state of our research.

The role of DSB in GPCR has also been investigated by others. For example, [29] describe decreased ligand binding after the removal of a DSB in the extracellular part of the MOR. A review article [30] mentions decreased agonist affinity at the CXC-chemokine receptor 4 and increased constitutive activity of the angiotensin II type 1 receptor after breaking extracellular DSBs. It must be kept in mind that ligands may cleave extracellular DSB in MOR [31]. If this also occurs inside the binding pocket, radical-induced DSB formation may not play a major role for opioid receptor activity.

In summary, comparing the influence of two prominent inflammatory mediators (pH and radicals) on ligand-induced opioid receptor function, it seems that pH has a higher impact than radicals under the chosen parameters. When designing novel opioid painkillers devoid of side effects elicited in non-injured environments, pH-sensitivity may be more important than radical-sensitivity. Given the high degree of homology between GPCRs [1], our current studies may be applicable to other signalling pathways (e.g. from receptor to nucleus [12]), to GPCR involved in other diseases (e.g. cancer, high blood pressure, addiction, depression, arthritis) or even to non-human GPCRs in deranged environments (e.g. in animals or plants exposed to ocean acidification).

## Author Contributions

Conceptualization: Stefanie Winkelmann, Marcus Weber, Christof Schütte, Christoph Stein. Formal analysis: Stefanie Winkelmann, Marcus Weber, Christoph Stein. Investigation: Arne Thies, Surav Ray, Vikram Sunkara, Hanna Wulkow, M. Özgür Celik, Fatih Yergöz. Supervision: Christof Schütte, Stefanie Winkelmann, Marcus Weber, Christoph Stein. Writing – original draft: all authors. Writing – review and editing: all authors.

## Acknowledgements

This research was partially funded by the Deutsche Forschungsgemeinschaft (DFG, German Research Foundation) under Germany’s Excellence Strategy – The Berlin Mathematics Research Center MATH+ (EXC-2046/1 project ID: 390685689), by DFG-STE 477/19-1, and via Project C03 of CRC 1114.

Codes and data are available at https://github.com/user3849/MOR1.

## A Supporting information

### A.1 MD simulations

For creating the different possible protonation states of the MOR amino acid residues in inflamed and in healthy tissue in the computational molecular model, a virtual “pH” value has to be selected. In inflamed tissue we selected “pH 5”, in healthy tissue “pH 7”. This only accounts for the modelling of protonated vs. deprotonated amino acids, because individual *H*^+^-ions are not part of the modelling. In reality, we always will find a mixture between different protonation states of amino acids. For example, at normal pH 7.4 there is also a small percentage of protonated histidines. Thus, in the following passages “pH 5” and “pH 7” just accounts for the parameter setting during the modelling step. Furthermore, the argument that there is always a mixture of different states also led us to take into account transient states with an additional DSB at pH 7 and without an additional DSB at pH 5. Systems at pH 5 and pH 7 without a ligand were also considered, both without and with an additional DSB, for comparison with systems where a ligand was present in the vicinity of the binding region.

For molecular modelling, the rat MOR structure was procured from the RCSB database (Protein Data Bank (PDB): 6DDF). Protonation states of the individual amino acid residues in the receptor were determined based on calculations at pH 5 and pH 7. The histidine imidazole side-chain has a pKa value of 5.97 [32]. Hence, these two levels of system acidity represent histidine states below (pH 5) and above (pH 7) the side-chain pKa. Other amino acids retain their protonation states as observed at normal pH (7.4). The protonated form of fentanyl, and the protonated and deprotonated forms of NFEPP [2] were sketched and parameterised using the CHARMM-GUI *Ligand Reader & Modeler* [33]. The protonated fentanyl was positioned onto the MOR at pH 7 with the Autodock program [34]. The docking calculations employed the Gasteiger-Marsili charges [35]. Autogrid was used for grid preparation, with grid spacing set to 0.65 Å to cover the entire receptor. Lamarckian genetic algorithm [36] was used to perform ten docking runs; with the rates of gene mutation and crossover kept at 0.02 and 0.8, respectively for the LUDI scoring function employed [37]. Remaining docking parameters were kept at their default values. The receptor-ligand complex with most energetically-favourable docking was used for further simulations. For similar starting conformations, the other ligands were aligned to the docked protonated fentanyl with the *RMSD Trajectory Tool* of VMD [38].

The receptor-ligand complexes were inserted into the 1-palmitoyl-2-oleoyl-sn glycerol-3-phosphatidyl choline (POPC) bilayer models using the CHAMM-GUI *Membrane Builder* [39]. Similar to [7], MD simulations were performed with GRO-MACS 2019.6 [40], using the CHARMM36m force-field for the ligands [41], receptor [42] and lipids [43]. The CHARMM TIP3P water model [44] was used as an explicit solvent. Sodium and chloride counterions were added to neutralise the excess charge and obtain a salt concentration of 0.15 M. The particle mesh Ewald (PME) method [45] was employed to calculate long-range Coulombic interactions, with a 1.2 nm cut-off for real-space interactions. A force-switch function was implemented for the Lennard-Jones interactions, with a smooth cut-off from 1.0 to 1.2 nm. The temperature was maintained at 310 K using the Nosé-Hoover thermostat [46, 47]. System pressure was kept at 1 bar with the Parrinello-Rahman barostat [48], using a semi-isotropic scheme where pressure along *x*-*y*-directions and the *z*-direction were coupled separately. Coupling constant and compressibility of the barostat were set to 5 ps and 4.5 × 10^−5^ bar, respectively. The LINCS algorithm [49] was used to constrain the covalent bonds between hydrogen and other heavy atoms, allowing a simulation time-step of 2 fs.

All simulation systems went through consecutive minimisation, equilibration and production runs, using the GROMACS scripts generated by the CHARMM-GUI [39]. First, the systems were energy minimised with steepest descent algorithms, followed by six-step equilibration runs. The first two runs were performed in the NVT (constant particle number, volume, and temperature) ensemble and the remaining runs in the NPT (constant particle number, pressure, and temperature) ensemble. Restraint forces were applied to the ligand, receptor, lipids, and water molecules, and *z*-axis positional restraints were placed on lipid atoms to restrict their motion along the *x*-*y*-plane. These restraints were progressively reduced during the equilibration process. Additional restraints were applied throughout equilibration to keep the distance between the crucial ASP 147^3.32^ and HIS 297^6.52^ residues of the MOR binding site [7] and the ligand molecule to the minimum possible. This ensured similar receptor-ligand starting conformations for the production runs of all the systems. Ultimately, unrestrained NPT production runs of 10 ns were performed, with periodic boundary conditions along all three orthonormal directions. Production run trajectories were saved every 10 ps, and processed with GROMACS analysis tools to generate the required information. VMD software was used for visualisation.

### A.2 In vitro experiments

#### Measurement of calcium currents in sensory neurons

To mimic the mechanisms underlying *in vivo* opioid analgesia, we examined calcium currents in sensory neurons harvested from rodents using a patch clamp protocol modified from [50]. The following chemicals were used: Dulbecco’s Modified Eagles Medium (DMEM)/HAM’s F-12 medium (Biochrom F4815, Berlin, Germany), Penicillin (10,000 U), Streptomycin (10 mg/ml), 1.25% Collagenase (Sigma-Aldrich C0130, Taufkirchen, Germany), 2.5% Trypsin (Sigma-Aldrich T0303), acridine orange/propidium iodide (Logos, Villeneuve, France), CaCl_2_· 6H_2_O, TEA-Cl_2_, 4-(2-hydroxyethyl)-1-piperazineethanesulfonic acid (HEPES), d-glucose, CsCl, MgCl_2_, ethylene glycol-bis-(*β*-aminoethyl ether)-N,N,N’,N’-tetraacetic acid (EGTA), Mg-ATP, GTP (Sigma-Aldrich).

Dorsal root ganglia (DRG) were harvested from naïve male Wistar rats (200-300 g; Janvier, Le Genest-Saint-Isle, France). Rats were killed by an overdose of isoflurane (AbbVie, Wiesbaden, Germany). The thoracic and lumbar spinal regions were exposed, DRG were collected in a digestive solution with 1.25% collagenase and incubated for 60 min at 37°C. After washing the cells three times with phosphate buffered saline (PBS), they were incubated in a digestive solution with trypsin for another 10 min at 37°C. After digestion, the tissue was triturated using plastic pipette tips and subsequently filtered through a 40 *µ*l filter. The filtrate was centrifuged, the supernatant was discarded and the pellet was resuspended in 1 ml culture medium (DMEM/HAM’s F12 supplemented with 1% penicillin/streptomycin and 10% horse serum). Cells were then seeded onto poly-L-lysine coated plastic culture dishes (35 mm) and placed in an incubator (5% CO_2_ at 37°C). One hour later, the cell cultures were topped up to a total of 2 ml of culture medium and cultured until patch clamp recordings, as previously described [51].

Recordings from DRG neurons were performed 24–48 h after plating. Cell viability was evaluated before the first experiment by an automated cell counter (Luna, Villeneuve, France) using acridine orange/propidium iodide. Recordings were carried out in whole-cell voltage clamp mode. After washing with PBS, cells were bathed in an extracellular buffer (ECS) (10 mM CaCl_2_ · 6H_2_O, 130 mM TEA-Cl_2_, 5 mM HEPES, 25 mM d-glucose; adjusted to pH 7.4 or 6.5; all from Sigma-Aldrich) and visualised using a Zeiss Axiovert 200 inverse microscope (Zeiss, Jena, Germany). Patch pipettes (resistance 3.5–8 MΩ) were produced from Borosilicate glass capillaries using a Sutter P-97 puller (Sutter Instruments, Novato, CA, USA) and filled with intracellular buffer (105 mM CsCl, 2.5 mM MgCl_2_, 40 mM HEPES, 10 mM EGTA, 2 mM Mg-ATP, 0.5 mM GTP, 5 mM d-glucose; adjusted to pH 7.4 or 6.5; all from Sigma-Aldrich). Currents were amplified and recorded using an EPC-10 patch amplifier and Pulse software (HEKA, Lambrecht, Germany). Extracellular buffer was added in a steady flow of 800–1,000 *µ*l/min using a pressurised application system (Perfusion Pressure Kit VPP-6; Warner Instruments, Hamden, CT, USA) and a suction pump. Opioid ligands (fentanyl, NFEEP, naloxone) were applied using a perfusion valve system (VC-6; Warner Instruments) to switch between vehicle (buffer) and the test compounds. After reaching the “giga-seal” at −60 mV, the membrane patch was breached to achieve whole-cell configuration. Only cells showing proper action potentials were selected for further experiments. The currents were initially recorded at a holding potential of −80 mV in ECS buffer in the absence of opioid ligands. Immediately thereafter, the cells were depolarised to +10 mV (100 ms) for eight times after 20 s intervals. During the first five cycles, only ECS was applied. On the sixth cycle, an opioid agonist (fentanyl, NFEPP) was added to the solution. During the last two cycles, the opioid antagonist naloxone was used to remove the agonist. All recordings were performed at room temperature.

#### Measurement of G-protein activation

Because these experiments require genetic alteration (by transfection) of cells, we performed these measurements in commonly used human embryonic kidney (HEK293) cells (RRID:CVCL 0045, German Collection of microorganisms and Cell Cultures, Braunschweig, Germany). All chemicals were purchased from Sigma-Aldrich (Taufkirchen, Germany), unless otherwise stated. [^35^S]-guanosine-5’-O-(3-thio)-triphosphate ([^35^S]-GTP*γ*S) was purchased from Perkin Elmer (Waltham, USA). Cell culture reagents were purchased from Biochrom (Berlin, Germany).

Cells were maintained in DMEM supplemented with fetal bovine serum (Biochrom), penicillin (100 U/ml, Biochrom) and streptomycin (100 *µ*g/ml, Biochrom) with or without geneticin (G418, 100 *µ*g/ml, Biochrom), in 5% CO_2_ at 37 °C as described before [2]. Cells were passaged 1:3 −1:20 every second to third day from p8 and p28 depending on confluence. Cells were plated on culture dishes coated with poly-L-lysine 24 h before transfection. 24 h after seeding, confluent cells (70-90%) were transfected with 1 *µ*g per 200 *µ*l transfection mix of each plasmid containing the different cDNAs using X-tremeGENE HP DNA Transfection Reagent (Roche, Mannheim, Germany) following the manufacturer’s instructions. For stable transfection, pcDNA™3.1+ carrying the rat MOR provided by Christian Zöllner (University Hamburg, Germany) was linearised with restriction enzyme Bg1II (NEB, Frankfurt, Germany), and linearisation was verified by agarose gel electrophoresis. After 48 h, the medium containing the transfection reagent was removed and replaced by complete DMEM with 10% fetal bovine serum and penicillin/streptomycin (100 U/ml). Successfully transfected cells were selected by adding G418 (500 *µ*g/ml) into medium that was renewed every 2 to 3 days. Monoclonal cell lines were then created 17 days post transfection by picking single colonies of stably transfected cells using a 100 *µ*l pipette and transferring them to poly-L-Lysine coated wells of a 96-well plate. Cells were grown to confluence and successively transferred to larger culture flasks in the continued presence of 500 *µ*g/ml G418. Antibiotic concentration was reduced to 100 *µ*g/ml when the cells were moved to 75 cm^2^ culture flasks. Monoclonal cell lines were further characterised based on immunocytochemistry, MOR mRNA expression, subjective impression of cell growth and overall cell morphology, as described previously [2]. Stably transfected cell lines were cultured for a maximum of 23 passages.

Protein concentrations were determined with the Bradford assay using Coomassie Brilliant Blue G-250 dye (Bio-Rad Laboratories GmbH, München, Germany) that shifts absorption from 465 to 595 nm upon binding to proteins. The relationship between measured absorbance and protein concentration was established based on a standard curve obtained from fixed protein solutions of known composition and concentration. These measurements were performed in duplicates using Bio-Rad Protein Assay Dye Reagent Concentrate with Bio-Rad Protein Assay Standard II (Bio-Rad). Samples with unknown concentrations, standards and dye reagent concentrate were diluted according to the manufacturer’s instructions, thoroughly mixed, and incubated for 5 min at room temperature. Absorption at 595 nm was measured in triplicates with a spectrophotometer. Generation of linear standard curves and interpolation of total protein concentration was performed by the device’s inbuilt software. A standard curve was generated for every experiment.

Membrane fractions were prepared from transfected HEK293 cells as described previously [52]. The cells were grown in 175 cm^2^ tissue culture flasks to approximately 90% confluence. Cells were then washed with Tris buffer (50 mM, Trizma preset crystals, pH 7.4; Sigma Aldrich), harvested with a scraper, homogenised using a mechanical disperser (Dispergierstation T8.10, IKA-Werke, Staufen, Germany) at maximum speed for 10 s and centrifuged at 42K×g for 20 min at 4°C (Avanti JXN-26 ultracentrifuge, Beckmann Coulter, Krefeld, Germany). Cellular pellets including membranes with embedded and anchored proteins were then resuspendend in Tris buffer for washing to separate them from cytosolic components by homogenisation and centrifugation at the same settings. Supernatants were discarded and the pellets were stored at −80 °C. On the day of usage, the pellets were thawed on ice in Tris buffer and homogenised. Total protein concentrations were determined as described above and homogenates were split according to the number of conditions tested in respective assay buffers.

The [^35^S]-GTP*γ*S binding assay was used to determine G-protein activation (as reflected by the exchange rate of GDP for GTP) at different H_2_O_2_ concentrations (0-1,000 *µ*M). GTP was replaced by a high concentration of [^35^S]-GTP*γ*S in the assay solvent, and the accumulation of [^35^S]-GTP*γ*S-bound G proteins in the membrane was measured. Membrane fractions were prepared with the following modifications: Membranes were homogenised and dissolved in HEM G-protein buffer containing 8 mM HEPES, 8 mM 4-(2-Hydroxyethyl)-1-piperazinepropanesulfonic acid (EPPS), 8 mM 2-(N-Morpholino)-ethanesulfonic acid (MES), 100 mM NaCl, 0.2 mM EGTA, 5 mM MgCl_2_ at pH 7.6, including freshly added 0.1% (w/v) bovine serum albumin (BSA). The desired amount of H_2_O_2_ was then added. To avoid interference with reactive oxygen species, the reducing agent dithiothreitol (DTT) (as originally used in [52]) was omitted. Basal [^35^S]-GTP*γ*S binding was assessed in the presence of vehicle without opioid ligands. In analogy to [53], 50 *µ*g of membrane fractions in duplicates were incubated with GDP (30 *µ*M) and [^35^S]-GTP*γ*S (0.05 nM) for 90 min at 30 °C. Unspecific [^35^S]-GTP*γ*S binding in the presence of non-radioactive GTP*γ*S (10 *µ*M) was subtracted to yield specific binding. Bound and free ligands were separated by rapid filtration under vacuum through Whatman GF/B glass fiber filters soaked in water followed by 6 washes with Tris Buffer. Bound radioactivity was determined by liquid scintillation spectrophotometry for ^35^S after overnight extraction of the filters in scintillation fluid optiphase HISAFE 3 (Perkin Elmer, Waltham, USA). Concentrations of radioactive compound were calculated based on the half life of ^35^S (87.4 days). Experiments were randomised to compensate for position effects in the filter apparatus or unequal sample processing times. Data processing and analysis were blinded for different H_2_O_2_ concentrations with the help of a colleague.

The data on dissociation of G-protein subunits, as measured by FRET, were extracted from [2]. Methodological details are described in [2].

#### Data Analysis

Experimental designs were randomised to compensate for the position effects on plates or filter apparatus and unequal sample processing time. Sample sizes were calculated using the G^∗^Power 3.1.2 program with *α* < 0.05, a power of 80% and a defined effect size (derived from pilot experiments). Analysis of concentration-response relationship was performed with simple linear regression using the GraphPad Prism 9 program (GraphPad, San Diego, USA) where y = [^35^S]-GTP*γ*S bound and x = [H_2_O_2_]. A P value ≤ 0.05 was considered statistically significant. Normal distribution of the data was assessed using the Kolmogorov-Smirnov test. Data are represented as means ± standard error of the mean (SEM).

#### Footnotes

1 In other papers like [15] this is referred to as switching the calcium channel from the “willing” state to the “reluctant” state.

